# Bundling Alleles Within Haplotype Blocks Improves QTL Detection Under Allelic Heterogeneity

**DOI:** 10.64898/2026.09.04.749340

**Authors:** Stan Oome, Mohammad Ghanbari

## Abstract

Allelic heterogeneity is a term from genetics which means that alleles that differ in primary sequence can have a similar phenotypic outcome. In other words, they are functional equivalents, and they naturally appear through convergent evolution under selection. Current GWAS has trouble detecting these instances, as allelic heterogeneity leads to signal dilution in these analyses, often leading to a LOD score that stays below detection thresholds. In this paper, we show a method that can overcome this problem by bundling haplotypes into Artificial Combined Markers. The created marker matrix can then easily be used in existing GWAS software.

## Introduction

Although substantial advances have been made in genetic and phenotypic technologies for studying living organisms, the overall framework of genetic association studies has remained largely unchanged. Recent progress has primarily focused on increasing the size and scale of genetic datasets and improving phenotypic measurements. Nevertheless, the same fundamental statistical framework—Genome-Wide Association Study (GWAS)—continues to be the dominant approach for identifying loci associated with traits (Korte & Farlow 2013).

The classic patterns of genetic inheritance were discovered by Gregor Mendel. For this, he used a biparental diploid system (pea plants), in which the now famous segregation ratios appeared. Plant breeding is, however, a relatively special genetic system, as (very) large biparental populations can be created (including self-pollinations), where modern limits are usually based on genotyping costs and phenotyping capacity. The availability of such populations has promoted the use of genetic map-based association methods, which can offer higher resolution than conventional GWAS (Luo et al., 2006).

Diploid biparental populations also have the advantage that Mendelian rules of genetics can be easily applied, such as segregation ratios, and the total amount of alleles per locus, since the original founding genetics of the population is completely known, and can often be deduced from the produced population (Strome et al., 2024).

In (many) animals, biparental populations are (much) more restricted in population size, especially from the female side. A more commonly used genetic system is a diversity panel, where diverse individuals are selected from the available genetic pool. And finally, we have natural populations, which can be used to study genetic variation in (wild) species where no selective breeding has been done, or in species where breeding experiments are deemed unethical, such as in humans. A key advantage of GWAS is its ability to utilize existing populations without the need to generate new mapping populations, which are costly, time-consuming, and sometimes unethical to develop (Yu et al., 2006).

In a diversity panel or natural population, when multiple genetic variants coexist, allelic heterogeneity can arise. Allelic heterogeneity is the genetic phenomenon where different mutations at the exact same gene locus result in the same or a highly similar phenotype. The resulting allelic pool exhibits frequency distributions that are often approximated by gamma or normal distributions. A gamma distribution is characterized by skewness for rare alleles and high density for common alleles, whereas a normal distribution concentrates alleles at intermediate frequencies. However, allelic diversity does not necessarily translate into phenotypic diversity, since genotypes are composed of combinations of alleles. This complexity can reduce the statistical power of single-marker approaches such as GWAS (Leroux et al., 2014).

In its simplest form, a GWAS can be viewed as a collection of independent tests in which each marker is evaluated for association with a phenotype. Although computationally efficient, this framework does not capture dependencies among alleles within genes. In particular, alleles with different DNA sequences but with similar biological functions may segregate together at varying frequencies, leading to signal dilution when molecular markers are not functionally specific. This is a common phenomenon because molecular markers are designed according to variation in DNA sequences. An intuitive loss-of-function example of allelic heterogeneity is a null-allele: these can differ in their way of not being able to perform the function, but can do so in many different ways based on primary sequence: premature stop codon, (much) lowered gene expression due to mutation(s) in the promotor, a changed amino acid that prevents the allele from functioning, alternative splicing, etc., each linked to different genetic markers on the same locus. A known gain-of-function example of allelic heterogeneity is lactase persistence, where the human species on several occasions independently evolved the capacity to digest lactose beyond infancy due to convergent evolution on the same locus (Gerbault et al, 2011).

From an evolutionary perspective, such dilution is expected. Mutation continuously generates new allelic variants from ancestral states, many of which do not produce measurable phenotypic differences. Consequently, independent testing of markers may fail to fully capture the functional architecture of underlying traits. This limitation contributes to the well-known problem of missing heritability, in which observed phenotypic variation cannot be fully explained by identified genetic variants (Manolio at al 2009). A portion of this gap may arise from the inability of standard marker-based approaches to represent functionally equivalent alleles.

To address this issue, genomic prediction methods have been developed. Genomic prediction relies on the collective contribution of many markers rather than on identifying a single marker with a large effect (Meuwissen et al., 2001). Although genetic relatedness among individuals can introduce false positives in GWAS, the same relationships are exploited in prediction models to infer phenotypic outcomes. This shared dependence on genomic similarity highlights the complementary nature of association and prediction approaches. Over the past decades, genomic prediction has become widely adopted in plant and animal breeding, accompanied by substantial methodological advances (Breen et al., 2022). However, identification of quantitative trait loci (QTLs) remains essential and is often more informative than prediction alone, particularly in the early stages of breeding programs.

The growing availability of multiallelic markers is transforming association studies, especially in (but not limited to) polyploid species. In these systems, multiple alleles can coexist at a single locus, making biallelic SNP-based representations less realistic and often less informative. Consequently, reliance on biallelic markers may reduce the power to detect QTLs. Several studies have proposed multiallelic marker frameworks and multiple-regression models, which often improve QTL detection compared with single-marker GWAS (Browning & Browning 2007, Leyva-Pérez et al., 2022, Vexler et al., 2026). However, these approaches typically require the construction of haplotype blocks from biallelic markers or rely on sequencing-based genotyping to directly define multiallelic loci.

In most existing frameworks, haplotype blocks are encoded into design matrices in which alleles are treated as predictors in mixed-effects or multiple-regression models. Results are reported as significance values and effect estimates for each allele or haplotype. While effective for prediction, these models provide limited insight into functional relationships among alleles. Moreover, estimated allele effects do not necessarily correspond to causal variants, as they are influenced by linkage structure, missing data patterns, and model specification (Navarro et al., 2022).

Machine-learning and multiple-regression approaches can further identify influential haplotypes and model interaction effects among alleles. However, their primary objective is predictive accuracy rather than elucidation of functionally coherent allele groups.

Associative heterogeneity has been widely applied in medical research to explore patterns of association between multi-allelic loci (SNP sets) and phenotypes as outcomes. A well-known example is the association of BRCA1 and BRCA2 alleles with similar effects on breast cancer risk (Woodward et al., 2022). In medical genetics, SNP-set–based approaches using GWAS summary statistics are commonly employed to study associative heterogeneity.

These methods generally fall into three categories:

1. Burden tests, which aggregate the effects of all SNPs within a haplotype block into a single score and test whether this score is associated with the phenotype (Lee et al., 2014, Pers et al., 2016).
2. Kernel-based association methods, which extend burden tests by assigning higher weights to rare variants and by squaring individual SNP effects to account for heterogeneous or opposing allelic effects (Wu et al. 2011, Ionita-Laza et al. 2013).
3. P-value combination methods, which aggregate SNP-level p-values and transform them into a single gene- or set-level p-value for testing association with the phenotype (Liu et al., 2020, Vsevolozhskaya et al., 2019).

Almost all SNP-set methods rely on summary statistics from single-marker GWAS analyses rather than individual-level genotype data. This is largely due to the massive number of individuals and markers involved, as well as the frequent unavailability of raw genotype data in many studies.

Here, we propose an extension of haplotype-based associative heterogeneity through a process termed “bundling”, which leverages individual-level data. After constructing haplotype blocks, alleles within each block are systematically combined to generate Artificial Composite Markers (ACMs). These ACMs represent higher-order aggregations of alleles and are analyzed as single variables using standard association testing frameworks.

### Mathematical Framework of Bundling

Consider a haplotype block (H) containing (k) distinct alleles,

(*H* = *a*_1_, *a*_2_, … , *a*_*k*_). For a population of (n) individuals, let (*x*_*ij*_) denote the dosage of allele (*a*_*j*_) in individual (i), where (*x*_*ij*_ ∈ 0,1,2) for diploid species and between 0 and *P* in polyploid species with ploidy level P.

The allelic composition of the haplotype block can therefore be represented by an (*n* × *k*) genotype matrix (X).

The bundling procedure begins by defining a subset of alleles (*B* ⊆ *H*), referred to as an ACM. It may contain any combination of alleles within a haplotype block.

Formula1 indicates that the total number of possible ACMs from the (k) alleles is equal to all possible combinations:

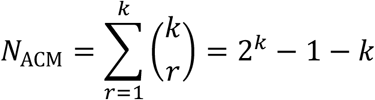

Thus, a haplotype block with 5 distinct alleles generates 26 possible composite markers (figure 1), whereas a block with 10 distinct alleles generates 1,013 possible composite markers. As can be seen, the number of ACMs increases exponentially, resulting in a substantial computational burden. Therefore, several filtering steps are applied to reduce the number of alleles before bundling. As a rule of thumb, alleles with a minor allele frequency below 0.01 are excluded. Second, redundant alleles with LD > 0.8 are removed. In addition, alleles with a frequency higher than 0.7 are excluded, because these alleles already have an optimal frequency for detection without bundling.

**Figure 1.**
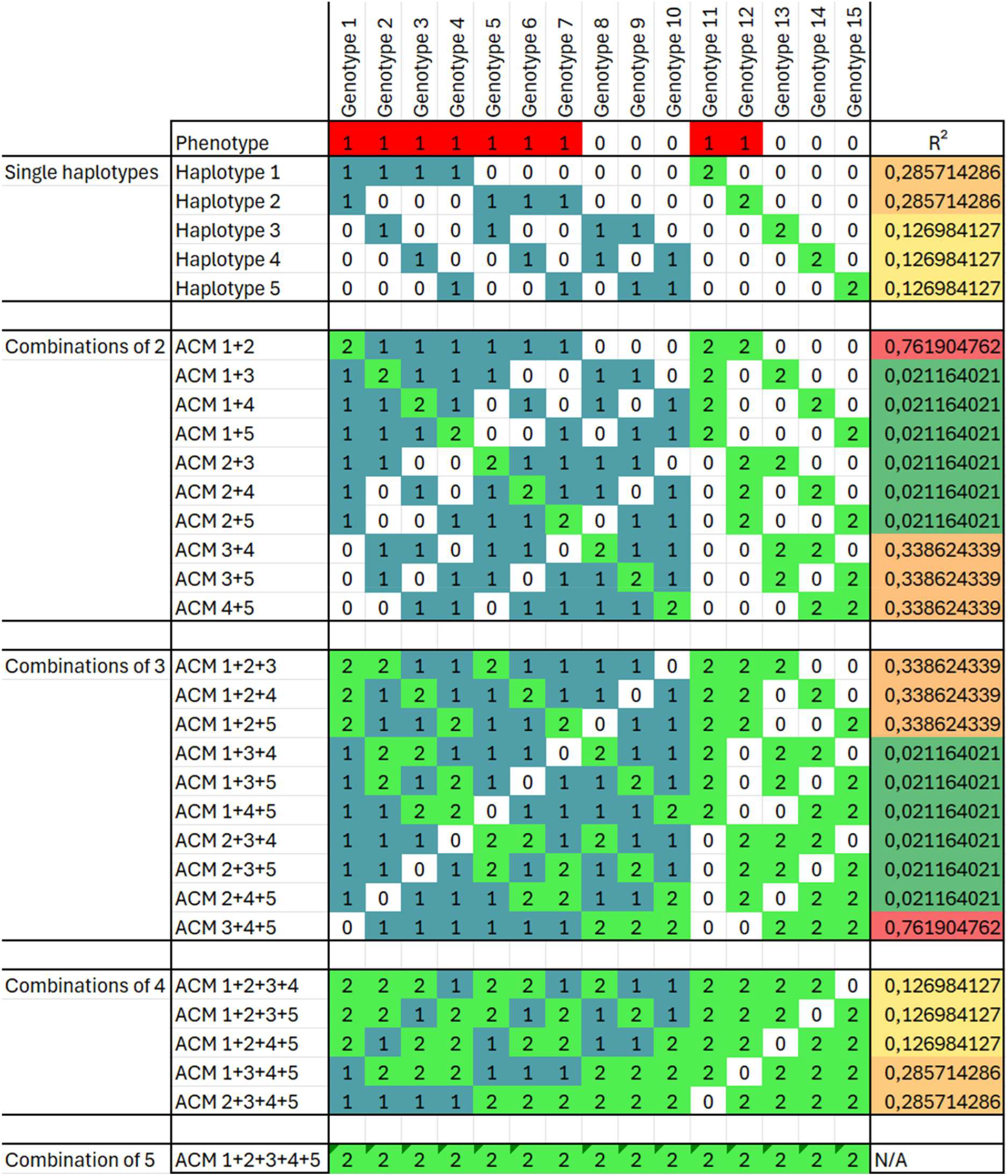
Example of a complete ACM matrix of a single locus of a genotyped diploid population of 15 individuals that contains 5 different haplotypes. 9 Individuals were phenotyped as having the trait “red”. None of the individual alleles can correctly explain the observed phenotype. After bundling of alleles, the combination of Haplotype 1 + Haplotype 2 most accurately matches the observed phenotype. This suggests haplotype 1 and haplotype 2 may be functionally interchangeable.

For each individual, a composite allele dosage is calculated as the sum of the dosages of all alleles belonging to the bundle:

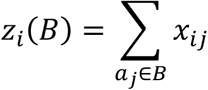

The resulting vector *z*_*i*_(*B*) represents an ACM that aggregates information across multiple alleles. Aggregated dosages may exceed the ploidy level in some genotypes, especially when bundling is applied to SNP markers. Therefore, ACMs with more than 10% of dosages above the ploidy level are regarded as artefacts and are therefore removed. In addition, in ACMs below this threshold, dosages exceeding the ploidy level are converted to the corresponding ploidy value in all ACMs. These filtering steps are applied to enable GWAS analyses in software that require a fixed ploidy level. However, packages such as GEMMA (Zhou 2012) and EMMAX (Kang et al., 2010) can handle such cases differently.

The ACM is subsequently treated as a conventional genetic marker and analyzed using standard association-testing procedures. For a quantitative trait (y), the association model can be expressed as

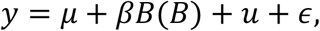

where (*µ*) is the intercept, (*βB*) is the effect of the bundle, (u) represents the random polygenic effect, and (ϵ) denotes the residual error (Yu et al., 2006). The statistical significance of the ACM is evaluated using the same test statistic applied in conventional GWAS.

This approach offers several advantages. First, it enables direct comparison between individual alleles and ACM groups in terms of statistical significance and effect size. Second, it can be implemented within existing GWAS pipelines without requiring specialized multivariate software, as ACMs are simply incorporated into standard genotype matrices.

In addition, bundling may increase the effective frequency of rare functional alleles. Variants that individually lack sufficient statistical power may become detectable when aggregated with functionally related alleles. As a result, the method may improve the identification of loci that are missed in conventional single-marker analyses.

Most importantly, bundling provides a framework for capturing the joint effects of alleles with shared or related biological functions. When multiple alleles correspond to similar functional states, their aggregation can strengthen association signals and increase estimated effect sizes. This directly addresses a key limitation of allelic heterogeneity, namely the dispersion of genetic signal across multiple functionally related variants.

A limitation of this approach is the increased number of markers generated through bundling, which also increases the multiple-testing burden. For large datasets, exhaustive analysis may exceed the computational capacity of standard computing environments. However, cloud computing platforms and distributed frameworks such as Apache Spark can make such analyses feasible at scale.

When such resources are unavailable, a two-stage strategy can be used. First, multiallelic association methods identify haplotype blocks with strong phenotypic effects (Zheng et al., 2015, Navarro et al., 2022). Bundling is then applied only within these regions, substantially reducing computational demand while retaining analytical power.

To evaluate the proposed method, we conducted simulation studies under four distinct genetic scenarios. A population of one thousand individuals was simulated at a locus with four alleles. For all simulated scenarios, an Ordinary Least Square method (OLS), as a standard GWAS, is run to test association between markers and the simulated phenotypes. A normal distribution with a mean of zero and a standard deviation of 0.5, which represents the measurement error, was added to the phenotypic value. For the sake of simplicity, the p value from OLS test is converted to LOD value using

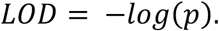

LOD of single and bundled markers (ACMs) are displayed as Manhattan plot and Bonferroni threshold according to number of tests that are used to determine the false discovery rate.

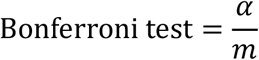

Where *α*= 0.05 and *m*= number of independent tests (markers or marker combinations) In the first scenario, the frequency distribution of alleles was 10% for two of the alleles and 1% for the other two. For phenotypic simulation, one allele with a high frequency and one allele with a low frequency were selected with an additive effect of 1, assigned on the phenotype. The total number of possible ACMs out of 4 distinct alleles according to formula 1 equals to 11. A standard GWAS runs over marker matrix to obtain p values for all the markers. Figure 2 presents the LOD scores obtained for all possible ACMs consisting of up to four alleles. The highest association signal was observed for bundle {1+3}, which achieved a LOD score of approximately 130 and corresponded exactly to the two causal alleles used in the simulation. Although the individual alleles were detectable, combining the common and rare causal alleles into a single ACM substantially increased the LOD score. These results demonstrate that allele bundling can improve QTL detection, particularly when causal variants include low-frequency alleles whose effects are difficult to capture using conventional single-marker GWAS approaches.

**Figure 2.**
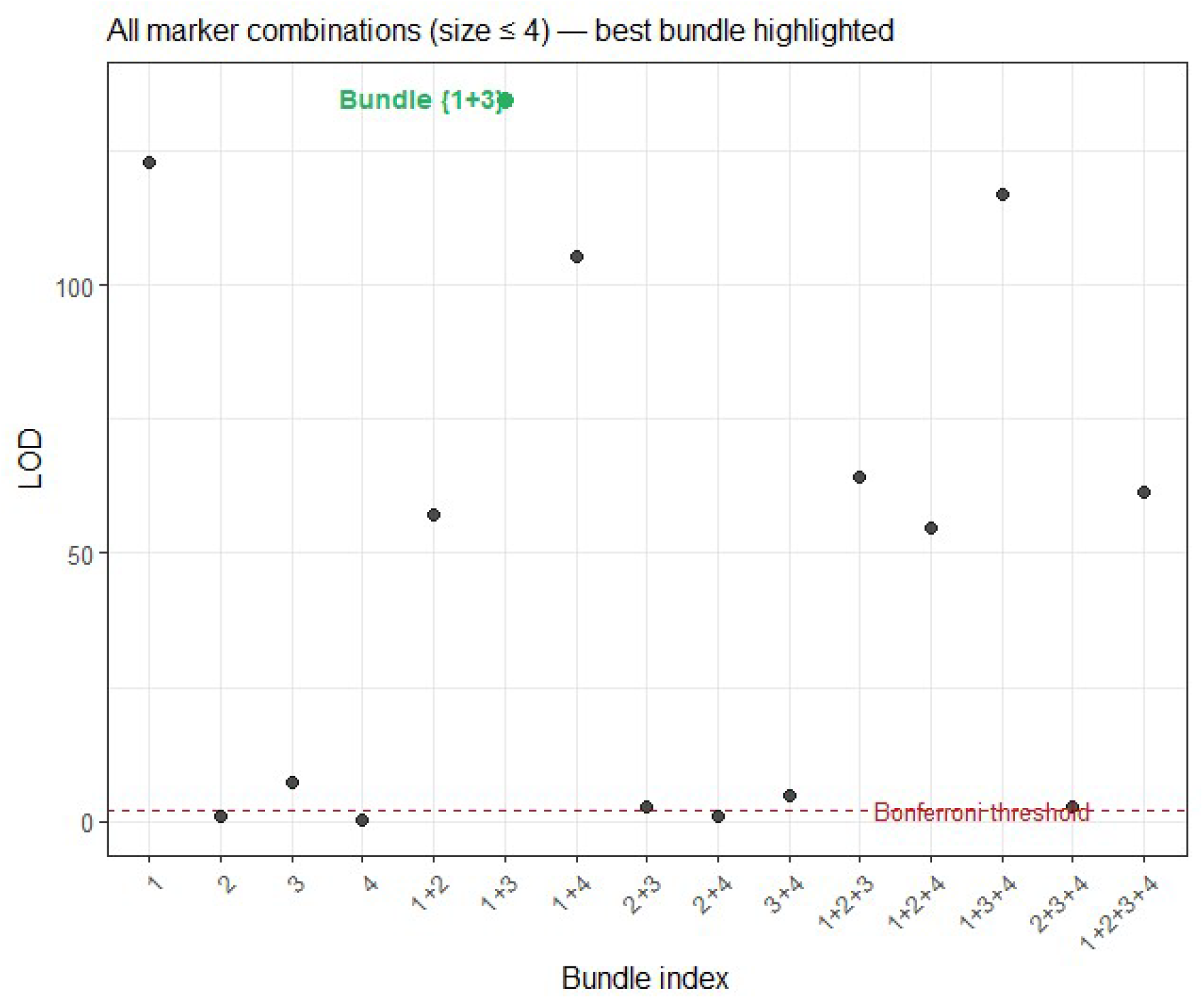
Manhattan plot of the first scenario demonstrates LOD scores for single markers and ACMs up to four markers. Marker 1 has an allele frequency of 10%, and marker 3 has an allele frequency of 1%. Additive effects of markers 1 and 3 are +1. The red dashed line denotes the Bonferroni-corrected significance threshold. The LOD score for ACM 1+3 is highlighted in green to indicate the detection power of bundling method in comparison with single GWAS.

In the second scenario, 4 alleles with an equal frequency of 0.25 were simulated. Alleles 1 and 3 are assigned to an additive QTL with positive effect of 1 and an additional environmental variance with mean 0 and standard error 0.5 are added to the genotypic effects. Results from the standard GWAS analysis are shown in figure 3. While individual allele markers exhibited moderate association signals, the ACMs corresponding to the bundle {1+3} produced the strongest association signal (LOD ≈ 300), substantially exceeding the significance threshold. This result demonstrates that ACMs can aggregate the effects of multiple causal alleles, thereby increasing the statistical power and improving QTL detection compared with standard single-marker GWAS analyses.

**Figure 3.**
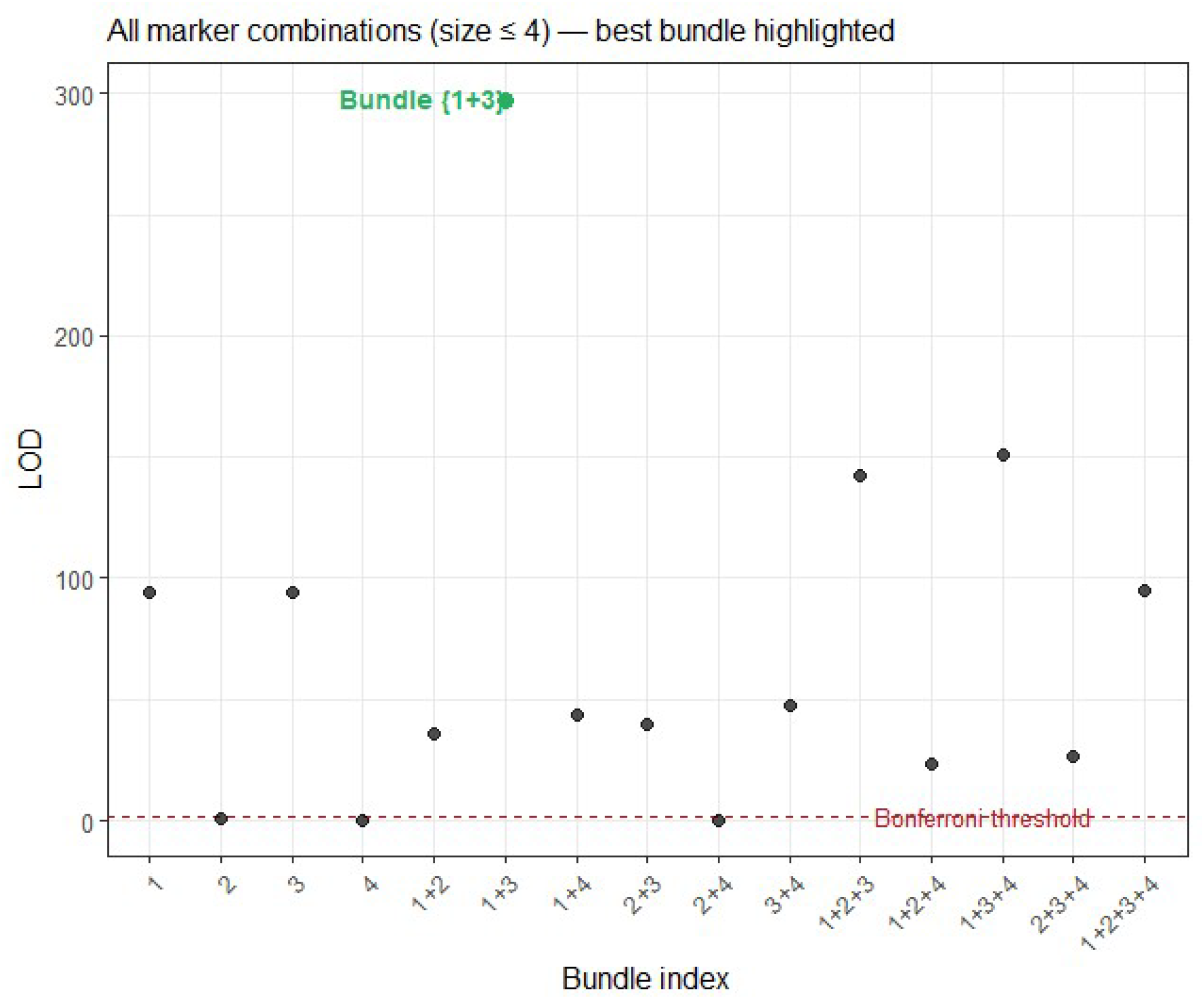
Manhattan plot of the second scenario demonstrates LOD scores for single markers and ACMs up to four markers. Marker 1 and marker 3 have an equal allele frequency of 25%. Additive effects of markers 1 and 3 are +1. The red dashed line denotes the Bonferroni-corrected significance threshold. The LOD score for ACM 1+3 is highlighted in green to indicate the detection power of bundling method in comparison with single GWAS.

The third scenario reflects a situation when alleles are carrying QTLs with opposites effects. There are 4 alleles with an equal frequency, but two alleles 1 and 3 are assigned to two QTLs with opposite effect of 1 and -1. The GWAS results in Figure 4 show that the individual allele markers corresponding to alleles 1 and 3 produced the strongest association signals, with LOD scores exceeding 100 and far above the Bonferroni significance threshold. In contrast, the ACM representing the bundle {1+3} produced a LOD score close to zero and was not significant. This occurs because of the positive effect of allele 1 and the negative effect of allele 3 cancel each other out when both alleles are combined into a single marker, effectively removing the underlying genetic signal.

**Figure 4.**
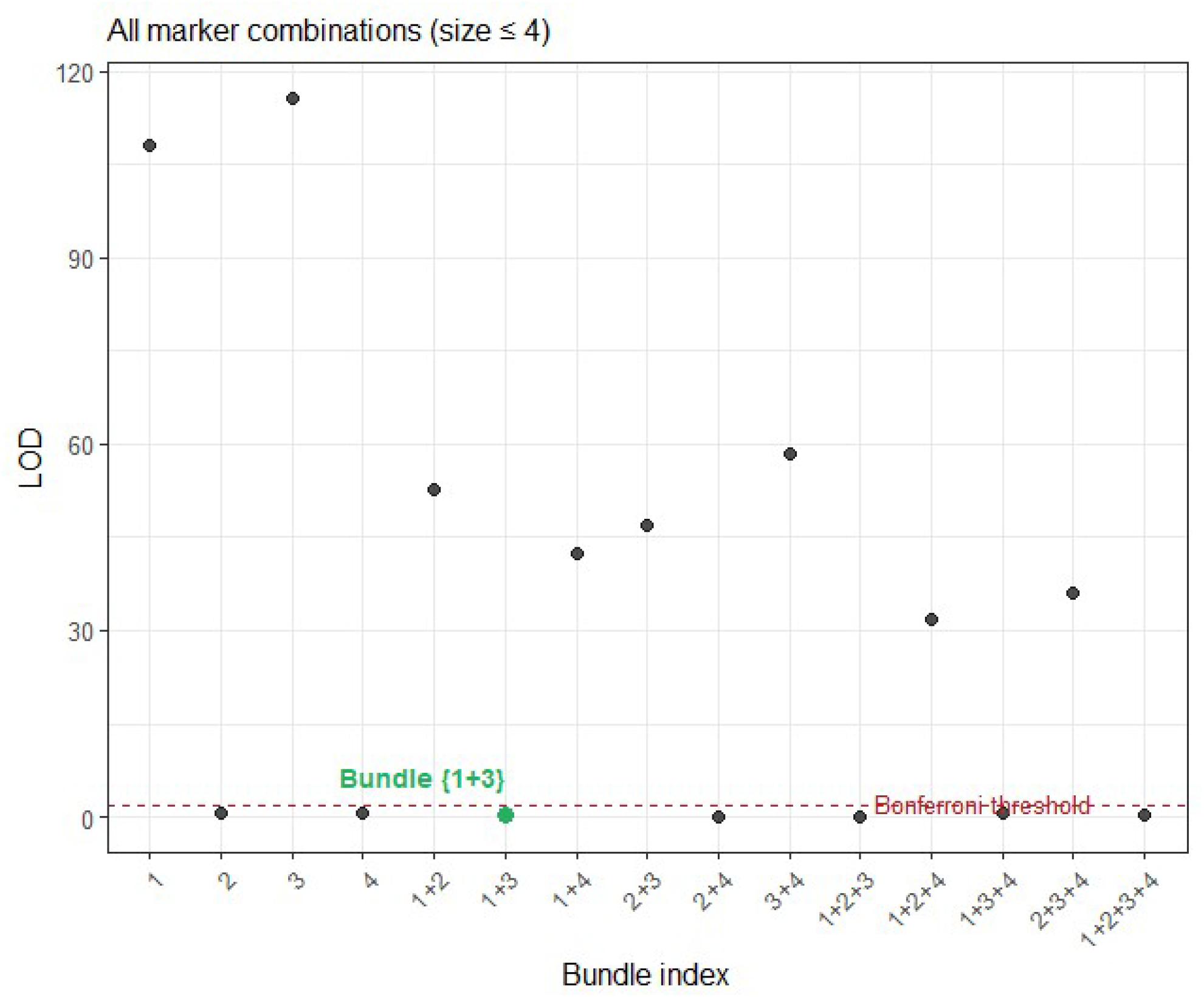
Manhattan plot of the third scenario demonstrates LOD scores for single markers and ACMs up to four markers. Marker 1 and marker 3 have an equal allele frequency of 25% but with the additive effect 1 for marker 1 and additive effect -1 for marker 3. The red dashed line denotes the Bonferroni-corrected significance threshold. The LOD score for ACM 1+3 is highlighted in green to indicate the detection power of bundling method in comparison with single GWAS.

In the fourth scenario (figure 5), the two QTL alleles with the same allele frequency of 0.25 have a different directional additive effect (i.e 1 and 0.3). This scenario reflects a more realistic genetic architecture in which multiple alleles contribute to the trait in the same direction but with different magnitudes.

**Figure 5.**
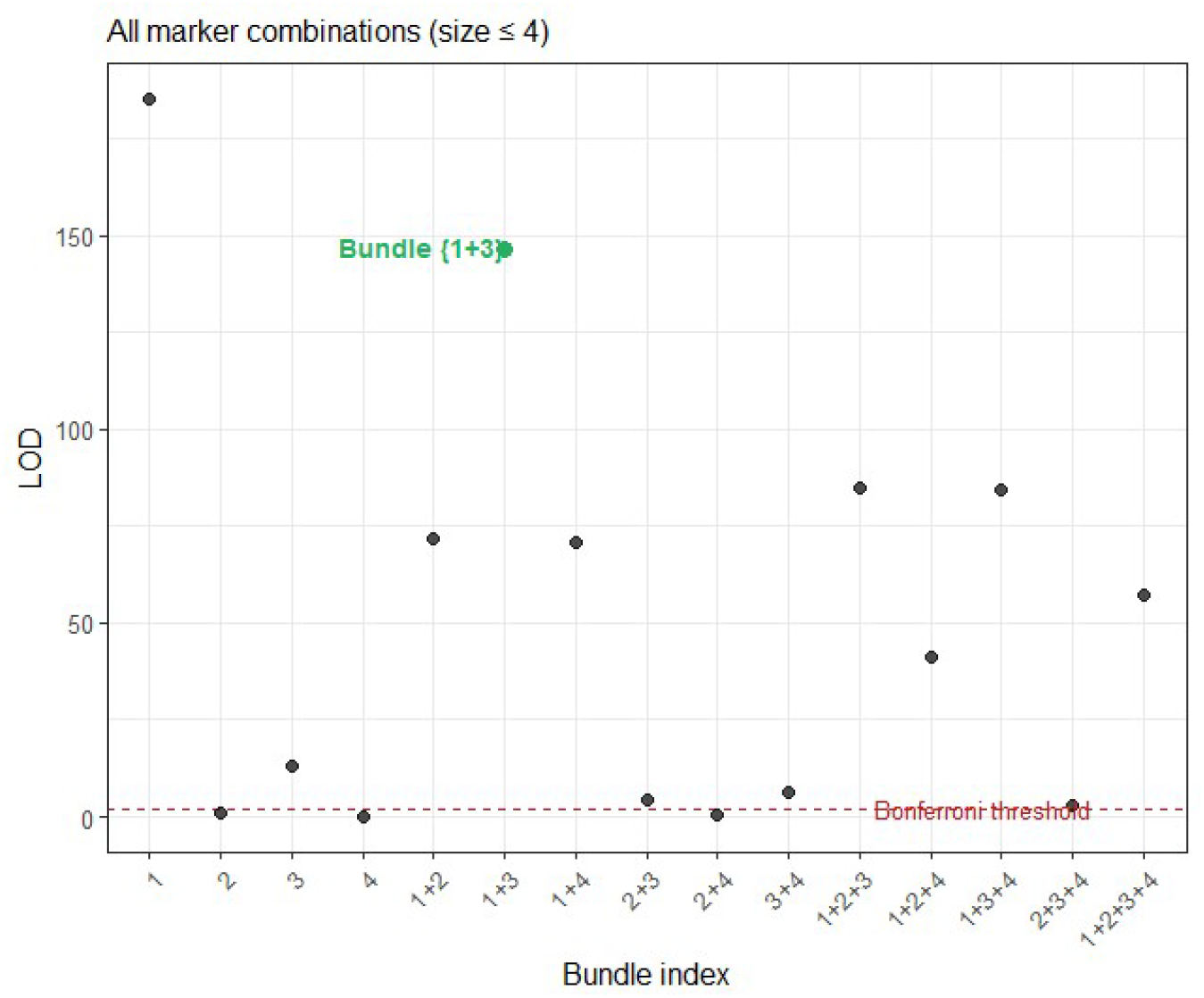
Manhattan plot of the fourth scenario demonstrates LOD scores for single markers and ACMs up to four markers. Marker 1 and marker 3 have an equal allele frequency of 25% but with the different directional additive effect 1 for marker 1 and +0.3 for marker 3. The red dashed line denotes the Bonferroni-corrected significance threshold. The LOD score for ACM 1+3 is highlighted in green to indicate the detection power of bundling method in comparison with single GWAS.

The results show that allele 1 generated the strongest individual association signal (LOD ≈ 180), consistent with its larger effect size. Allele 3, which had a weaker effect, produced a much smaller LOD score (LOD ≈ 13) but remained detectable. The ACM corresponding to the bundle {1+3} achieved a LOD score of approximately 145, greatly exceeding the significance threshold and representing the strongest signal among the allele combinations. This demonstrates that combining alleles with effects in the same direction can substantially increase detection power, even when the magnitudes of the effects differ.

However, the LOD score of the {1+3} bundle was lower than the signal observed for allele 1 alone. In the present scenario, the weaker effect of allele 3 contributed relatively little additional genetic variance to the bundle, limiting the gain achieved through aggregation. Nevertheless, the ACM still successfully identified the two causal alleles as belonging to the same functional group and produced a stronger signal than allele 3 alone.

## Discussion

The simulation results demonstrate that ACM-based GWAS can outperform conventional single-marker GWAS when a QTL is determined by multiple haplotypes with effects in the same direction. By combining independent haplotype effects into a single test, ACMs improve the signal-to-noise ratio, resulting in stronger statistical evidence for QTL detection and more accurate identification of the underlying genetic architecture, although the magnitude of improvement depends on the relative effect sizes of the alleles being combined. The greatest benefit is obtained when the bundled haplotypes contribute substantial and concordant effects to the trait. In scenarios 1 and 4 described here, unbundled haplotype 3 with a smaller effect size was already detected above the threshold individually. In real-world datasets, where additional sources of noise are often present, smaller-effect haplotypes may only become significant when bundled with other haplotypes that contribute effects in the same direction, further increasing the practical advantage of ACM-based GWAS.

The term “allelic heterogeneity” appears to be a *contradictio in terminis*, as an allele has to differ from another allele by definition. This apparent contradiction is due to the fact that the geneticist‘s and the molecular biologist‘s definition of an allele differ. The geneticist definition is “*One of two or more alternative forms of a gene that arise by mutation and are found at the same place (locus) on a chromosome, which are defined by their outcome—whether it is dominant, recessive, or creates a specific trait*”. For a molecular biologist, the definition of an allele is “*Any variation in the DNA sequence—a single base pair (SNP) or a segment of bases—at a specific genomic location*”. Two alleles might have different DNA sequences but produce the exact same functional protein (neutral variation), yet a molecular biologist would count these as different alleles, whereas a classical geneticist might group them together as a single allele. This distinction seems trivial, but any technique that tries to correlate genotype to phenotype runs into trouble, as “phenotype” refers to the genetic definition of an allele, as an outcome of the gene is measured, while “genotype” refers to the molecular definition, as genotyping works by identifying differences in primary sequence. Therefore, “allelic heterogeneity” as such, misses the point for a molecular biologist, and these instances would more likely be referred to as “functionally equivalent haplotypes”.

In this paper we show that bundling of haplotype blocks into Artificial Composite Markers (ACMs) may overcome this problem by uniting the definitions, which allows for standard GWAS calculations to identify haplotypes with a similar effect on a phenotype (the geneticists‘ allele), which normally may stay hidden due to signal dilution. For this phenomenon, we propose the term “Equistasis”, “Equi” meaning “the same” or “equal to”. “Equistatic” haplotypes are located at the same genomic location and differ in primary sequence but are functionally similar, and therefore biological/phenotypical equivalents of each other, regardless of how this equivalence is achieved (figure 6). When identified through ACM-based GWAS, it may be a case-by-case study to figure out by which exact mechanisms these particular haplotypes are equistatic.

**Figure 6.**
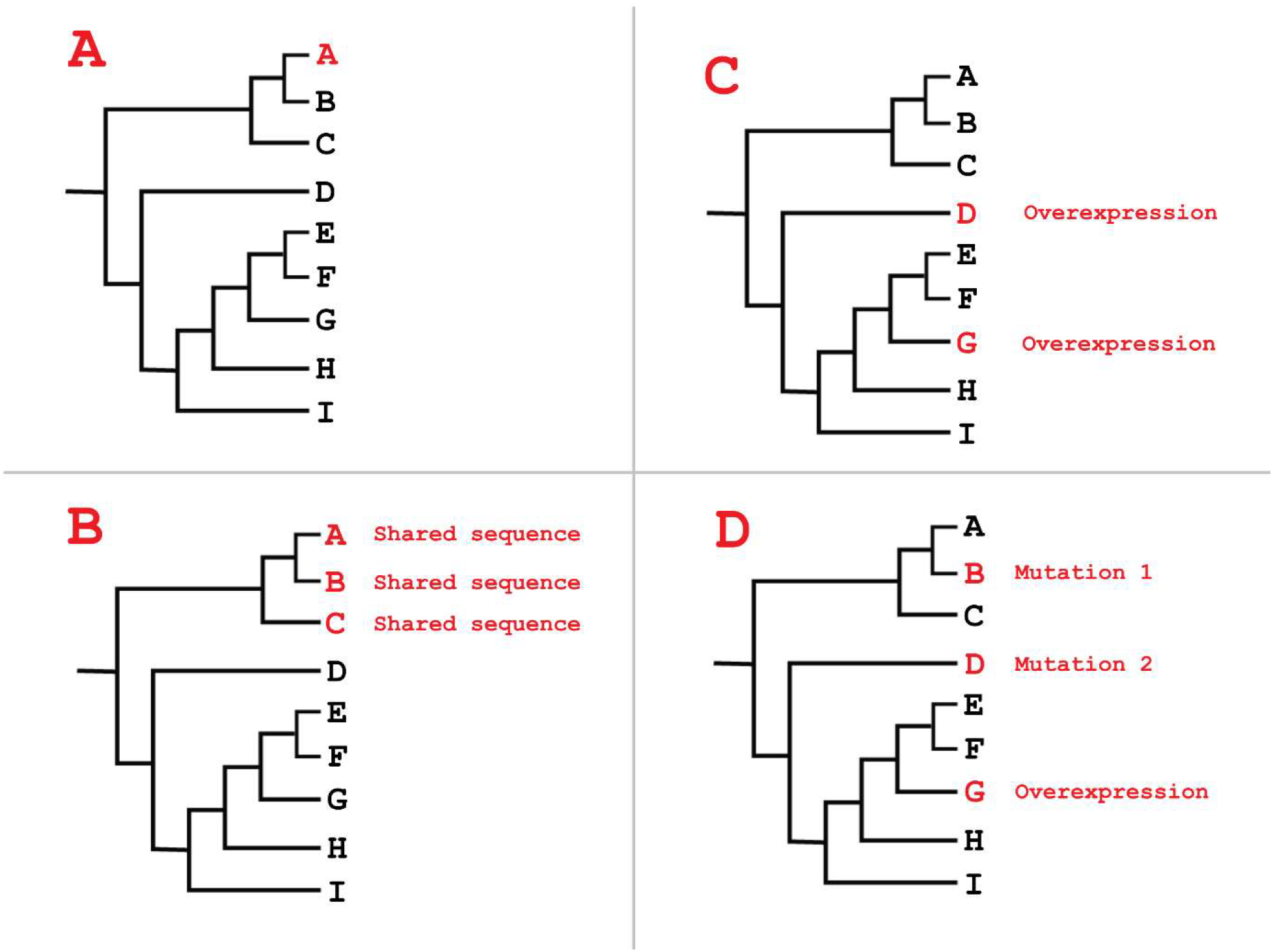
Different scenarios among 9 haplotypes in a population. **A:** Haplotype A correlates to a phenotype, suggesting haplotype A is functionally different from the other 8 haplotypes. This situation is generally detected by standard GWAS. **B:** Haplotypes A, B, and C all correlate to a phenotype. These can be identified by finding a correlation between a SNP that occurs on both these three haplotypes, but not in the other 6. The causal mutation is likely the same in all three haplotypes. **C:** Two haplotypes are overexpressors of the gene, and overexpression of the gene (technically a phenotype by itself) is correlated to the observed phenotype. From expression data it can be deduced that both haplotypes are independent overexpressors. It is quite possible that the observed overexpression is not caused by an identical mutation. **D:** There are multiple, independently evolved haplotypes, which all exercise a similar effect on the phenotype through a different mechanism, but which cannot be directly deduced by shared sequence similarity or expression level.

Applying this methodology to identify equistatic haplotypes in (existing or new) genetic datasets will probably lead to many new QTLs being identified and may clear up part of the “missing heritability” (Manolio et al., 2009). Another factor in missing heritability is epistasis, which is the effect of one gene being influenced, or masked, by one or more other genes. The matrix of ACMs created by our methodology can likely also be used to enhance epistatic analyses such as 2D epistatic Next-Gen GWAS (Carré at al 2024), although it may push the number of possible combinations outside the reach of currently practical calculation limits. Also, analyses based on the Hardy-Weinberg theorem may improve from using ACMs as input, as selection occurs on the geneticists‘ definition of an allele, not the molecular biologist‘s definition.

A difficulty in applying the construction of a good enough ACM matrix may be found in accurately covering all available alleles. Underestimation may occur due to lack of marker representation of (rare) haplotypes, while overestimation may occur due to the fact that not all SNPs are haplotype-specific (tag) SNPs. Overestimation can be identified by an ACM overshooting the ploidy state of the organism under investigation, but underestimations may be more difficult to detect, and may require imputation. The rise of technologies that produce microhaplotypes instead of single-SNP detection may be a factor that could alleviate this situation somewhat.

In conclusion, haplotype bundling assumes that multiple haplotypes within a locus may represent similar or partially equivalent functional states. When such haplotypes are tested individually, their effects may be distributed across multiple association signals, reducing statistical power. By aggregating haplotypes into Artificial Composite Markers, bundling compresses the allelic representation of a locus and concentrates the genetic signal into a lower-dimensional feature. Consequently, the method is expected to improve the detection of loci exhibiting allelic heterogeneity while remaining fully compatible with existing GWAS software, statistical frameworks, and molecular biologists‘ intuition in interpreting the results.

---

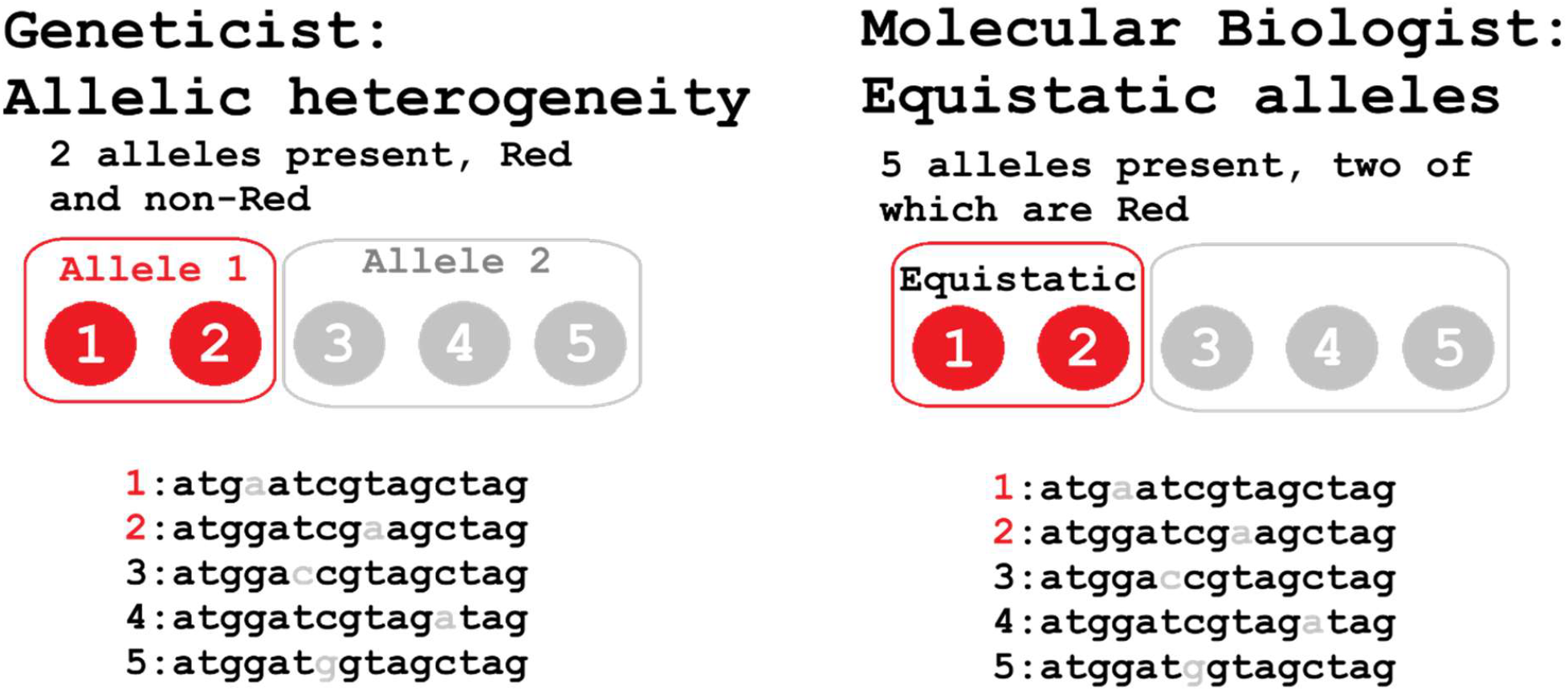

## Notes

### Competing Interest Statement

The authors have declared no competing interest.

